# Phenotypic variation and weak behavioral isolation in a putative hybrid zone between Andean *Atlapetes* brushfinches (Passerellidae)

**DOI:** 10.1101/2024.11.17.623919

**Authors:** Gabriela Muñoz-Quintana, Andrés M. Cuervo

## Abstract

Individuals in the contact zone between hybridizing species often exhibit intermediate phenotypes that vary smoothly in a clinal fashion from one species to the other. However, many tropical hybrid zones remain poorly documented due to limited specimens, small species ranges, and logistical challenges of accessing key localities within contact zones. We investigated phenotypic variation and species recognition between two endemic Andean brushfinches that replace each other geographically along the Central Cordillera of the Colombian Andes: *Atlapetes fuscoolivaceus* (Dusky-headed Brushfinch) and *A. flaviceps* (Yellow-headed Brushfinch). We scored phenotypic traits from 111 georeferenced photographs of live individuals and 40 museum specimens. Using cline-fit analyses for two hybrid indexes (face color pattern and overall body color), we tested whether phenotypic transitions were abrupt (suggesting a barrier, or a sharp replacement) or gradual (indicating hybridization) across their contact zone. We also conducted playback experiments to assess song discrimination as an indication of potential prezygotic reproductive barrier. Plumage traits varied clinally, with a center near Inzá, Cauca (2.55°N), and with notably different cline widths between facial (12 km) and body-color traits (73 km). This difference likely reflects greater sampling density for facial traits and possibly stronger selection on face coloration. Territorial *A. flaviceps* individuals responded to songs of both species (no difference in latency); however, response duration was longer when exposed to local songs, indicating weak behavioral isolation. Our results suggest that these two ecologically equivalent brushfinches maintain a hybrid zone where their ranges meet, with a smooth transition from a pure phenotypic form to the other, which is likely facilitated by weak prezygotic behavioral barriers to reproduction. This study represents the initial step in investigating phenotypic variation, speciation genomics, behavior and interaction at the hybrid zone between *Atlapetes* brushfinches in the Colombian Andes.

## INTRODUCTION

As allopatric sister species continue to diverge following the establishment of extrinsic barriers to reproduction, additional mechanisms may arise to reinforce or perpetuate isolation (Foster and Endler 1999; Coyne and Orr 2004; Edwards et al. 2005). However, if insufficient time has elapsed since their separation, behavioral and ecological prezygotic barriers that hinder interbreeding may not yet be fully developed (Mayr 1999; Coyne and Orr 2004; Price 2008). Given the critical role of song in courtship and mating for birds, vocal behavior acts as an essential mechanism of species discrimination, serving as a prezygotic barrier (Edwards et al. 2005; Price 2008; Uy et al. 2018; Lipshutz et al. 2019). Even closely related bird species can rapidly evolve differences in songs in allopatry (Podos and Warren 2007; Wilkins et al. 2013), particularly song learners like oscine passerines (Lachlan and Servedio 2004). However, if song signals remain similar and birds cannot discriminate vocalizations between species, other mechanisms such as reproductive asynchrony or postzygotic barriers may maintain isolation (Coyne and Orr 2004; Price 2008). Otherwise, incomplete reproductive isolation may lead to the formation of hybrid zones (Barton 1979; Barton and Hewitt 1985; Harrison 1993; Barton 2001; Abbott et al. 2013).

Hybridization is widespread in birds (Grant and Grant 1992; Ottenburghs 2023), occurring more frequently between lineages of rapid radiations or between populations at early stages of speciation (Barton 2001; Seehausen 2004; Price 2008). However, the number of stable hybrid zones, where introgression between two species is maintained at their range boundaries, is likely underestimated (Seehausen 2004; Price 2008; Ottenburghs 2023). Although hybrid zone biology is in its infancy in the Neotropics (Pulido-Santacruz et al. 2018; Lipshutz et al. 2019; Del-Rio et al. 2022), a few cases are known in Colombia (Haffer 1967; Morales-Rozo et al. 2017; Céspedes-Arias et al. 2021; Morales et al. 2021). For example, two *Myioborus* warblers hybridize where their ranges meet in southern Colombia and northern Ecuador (Céspedes-Arias et al. 2021; Cuervo and Céspedes-Arias 2023). Many more untold hybrid zones might be present in the rich evolutionary history of the Northern Andes.

A potentially undocumented hybrid zone may exist between species pairs in the genus *Atlapetes*, which have diversified recently in the montane forests of the Andes (Klicka et al. 2014; Sánchez-González et al. 2015) and show a tendency to hybridize (Fjeldså and Krabbe 1990; McCarthy 2006; Carantón-Ayala et al. 2018). Along the eastern slope of the Central Cordillera of the Colombian Andes, two *Atlapetes* brushfinches appear to replace each other (Fig. 1), but range limits are not clearly known (García-Moreno and Fjeldsa 1999; Ayerbe Quiñones 2023). *Atlapetes flaviceps* (Yellow-headed Brushfinch) occurs in the northern part of that range, from Antioquia (López-Ordóñez et al. 2013; Chaparro-Herrera et al. 2020) to approximately north-central Tolima (this study), whereas *A. fuscoolivaceus* (Dusky-headed Brushfinch) inhabits mainly in the southern end of the eastern Central Cordillera (this study) and the Upper Magdalena valley in southern Huila.

**Figure 1.**
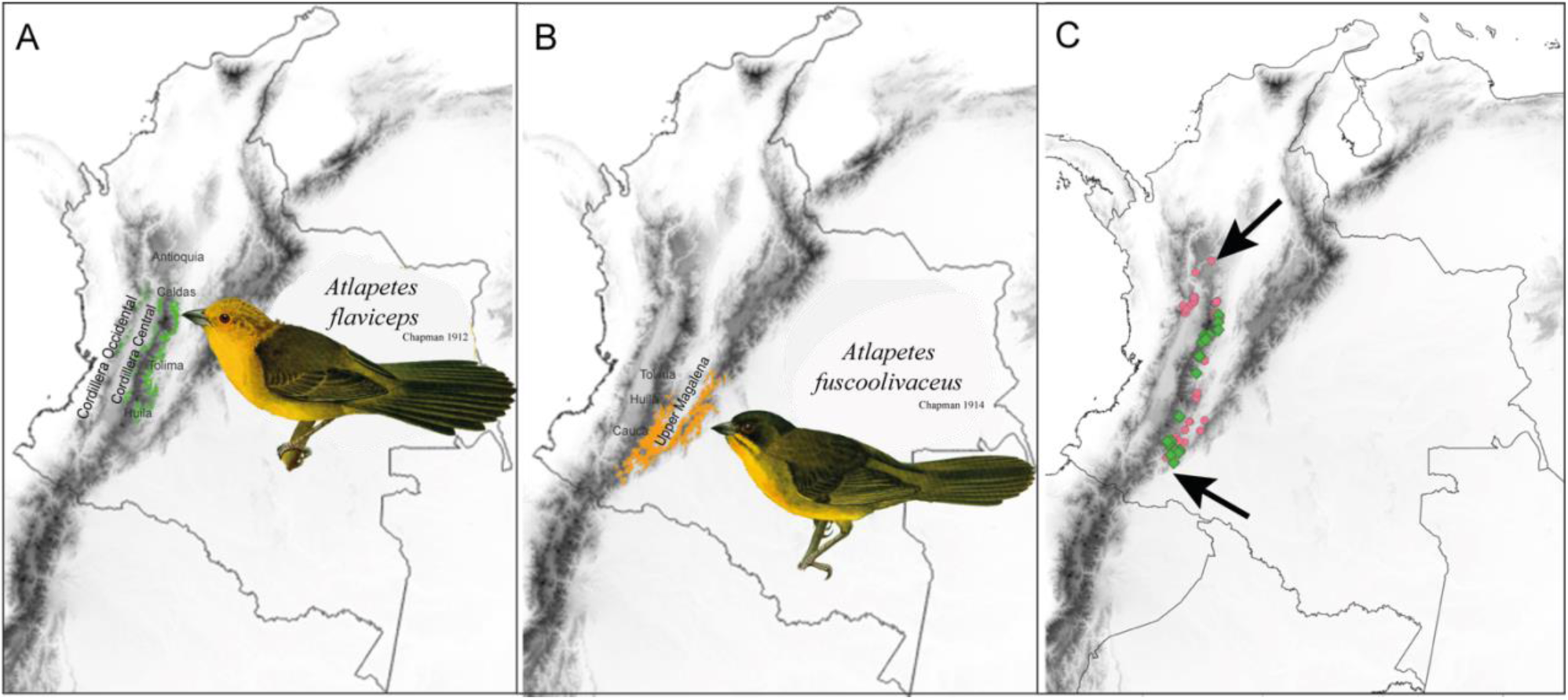
Approximate geographic distribution of **A**) *Atlapetes flaviceps* (Yellow-headed Brushfinch), **B**) *Atlapetes fuscoolivaceus* (Dusky-headed Brushfinch). Inset illustrations depict the main phenotypic characteristics of typical individuals in both species. Illustrations by Louis Agassiz Fuertes (in Chapman 1917) via the Biodiversity Heritage Library (https://www.biodiversitylibrary.org/item/117614). **C**) Geographic location of specimens (in green) and photographs (in pink) included for the inference of facial and body color clines in this study. The two arrows indicate the northernmost (*A. flaviceps*) and southernmost (*A. fuscoolivaceus*) data location within the study zone.

We noticed that several specimens do not perfectly match the stereotypical color patterns described for either species (Chapman 1912, 1914), especially those from southern Tolima, northern Huila, and eastern Cauca departments. Because these regions have been historically affected by poor road infrastructure, violence and social unrest, species diversity remains insufficiently analyzed (Olivares 1960; Ridgely and Gaulin 1980; Krabbe et al. 2005), leaving species diversity and range boundaries largely documented (Cuervo et al. 2005). For instance, although published sources suggest *A. flaviceps* occurs throughout southern Tolima, Huila and Cauca (Molina-Martínez 2014; Botero-Delgadillo et al. 2021), these areas have remained largely unsurveyed, and therefore, unconfirmed or misleading records of *A. flaviceps* (Ridgely and Gaulin 1980; Dunning 1982) have led to overestimation of its range (Botero-Delgadillo et al. 2022).

The gradation in measurable characters observed from the range of one species to the other and across the hybrid zone form a cline (Huxley 1938; Slatkin 1973; Endler 1977). Cline fit analyses allow describing the geographic transition of intermediate traits that arise from secondary contact between species (Mayr 1963; Endler 1977; Barton and Hewitt 1985; Curry 2015). Here, we aim to describe the color variation from one species to another and evaluate if there is any interspecies recognition. First, we used photographs and specimens to characterize the putative hybrid zone based on plumage coloration. Second, we conducted playback experiments, whose main goal was to evaluate the response of the local species (*A. flaviceps)* to its song, the foreign species song (*A. fuscoolivaceus*), and a negative control species song (*Zonotrichia capensis*). By studying geographic variation of color patterns using cline fit analyses and playback experiments, we expect to better understand potential behavioral barriers between the two species and the characteristics of the putative hybrid zone.

## METHODS

### Study species

The northern species *Atlapetes flaviceps* (Yellow-headed Brushfinch) is characterized by bright yellow face, crown and underparts, lacking any face marks, and its wings and upperparts are dark olive green (Chapman 1912). In turn, the southern species *Atlapetes fuscoolivaceus* (Dusky-headed Brushfinch) is covered with olive black in the face, crown and upperparts, whereas its underparts, including chin and throat, are olive-yellow, including prominent dark malar lines (Chapman 1914). Body size, behavior, elevational range, and habitat preferences are essentially the same in both species. The main phenotypic differences between “pure” individuals relate to facial color pattern, presence of malar line, and the tone of the yellowish underparts (brighter in *A. flaviceps*), and the dark olive upperparts (darker in *A. fuscoolivaceus*). Both species have small geographic ranges in the Colombian Andes (Fig. 1).

### Hybrid index scoring

To assess color pattern variation across the putative hybrid zone, we described plumage color variation from museum specimens and photographs. We developed a facial hybrid index using categories based on four facial patches scored from 40 adult specimens and 111 wild individuals photographed and obtained from public databases (Supplementary Table S1 and S2). These patches included the forehead, lores, ear patch, and malar lines, which show the greatest differentiation between “pure” individuals and exhibit polymorphism in the contact zone. For example, while *A. fuscoolivaceus* has distinct dusky malar lines, *A. flaviceps* has plain bright yellow throats without malar lines. Because several individuals showed intermediate phenotypes, we created intermediate value categories (Table 1).

**Table 1.**
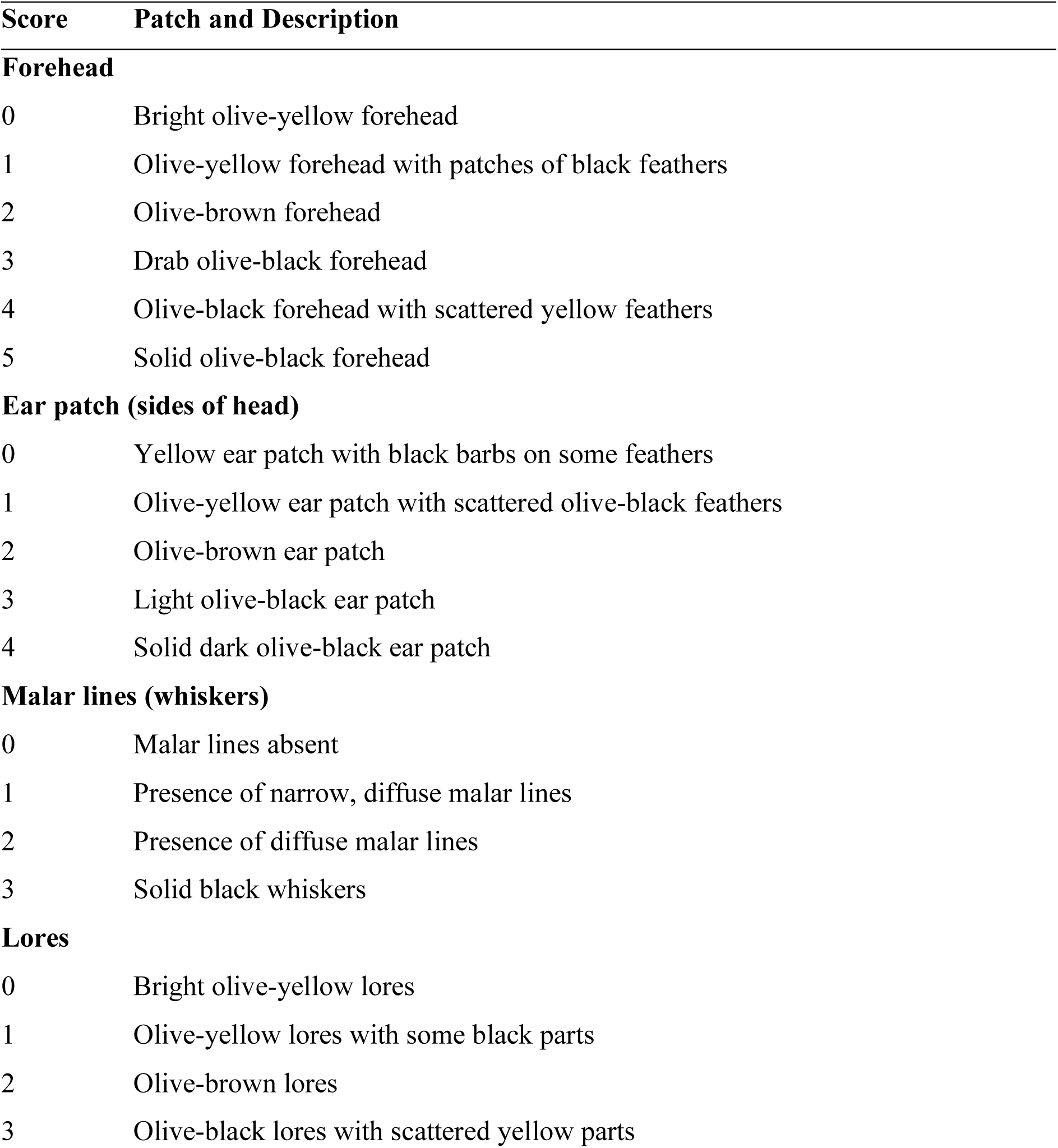
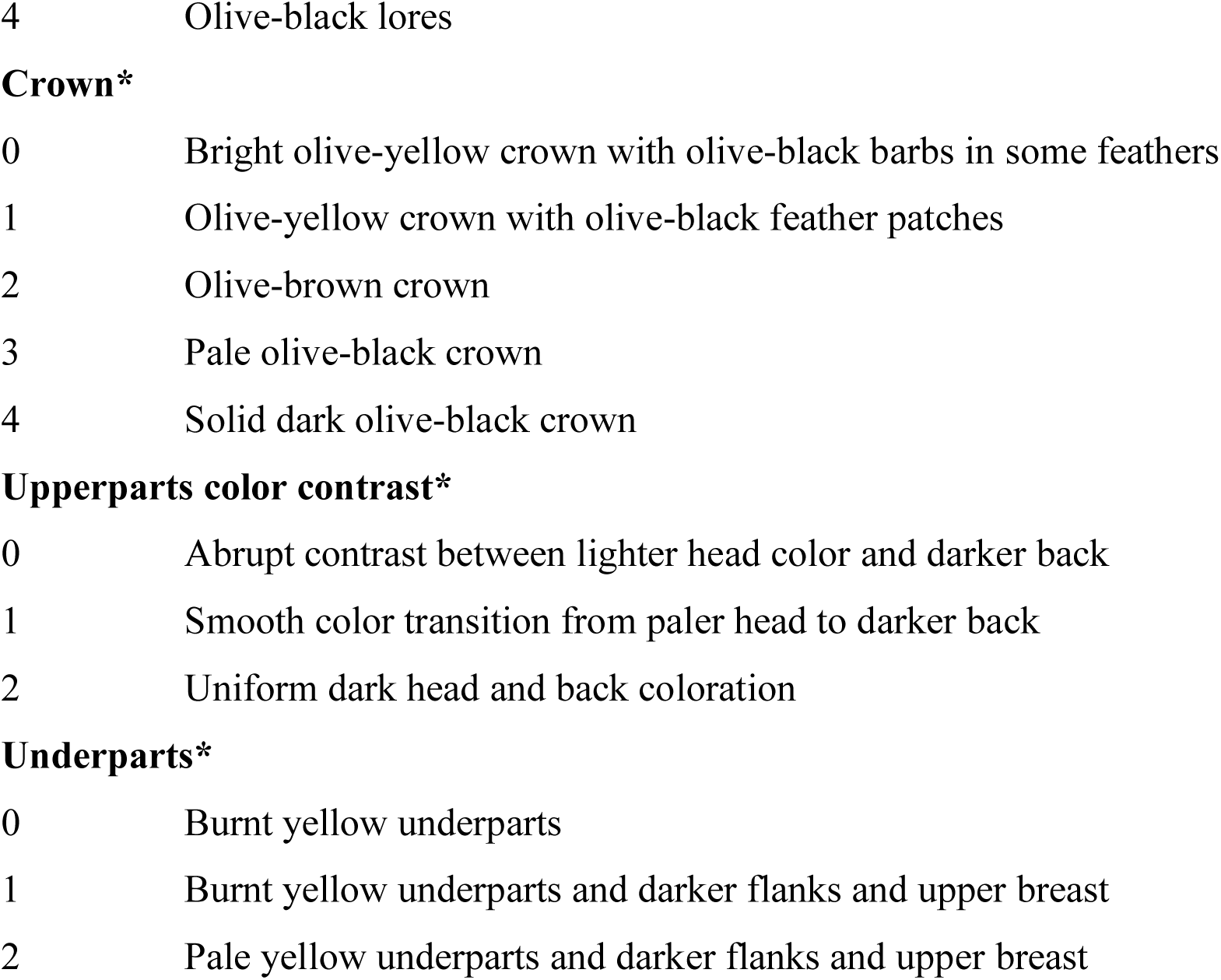
Phenotypic scoring system used to calculate facial and body hybrid indexes in *Atlapetes* brushfinches. Scores range from 0 (typical *A. flaviceps* phenotype with lighter coloration) to 1 (typical *A. fuscoolivaceus* phenotype with darker coloration). Upper facial patches (crown, lores, auriculars, and malar) were scored from both photographs (n = 111) and museum specimens (n = 40). Body patches (back, flanks, and undertail coverts) were scored from museum specimens only.

We then developed a body color hybrid index using only the 40 adult specimens. Following the same procedure as for the facial hybrid index, we scored three additional patches: crown color, upperparts color (back and rump combined), and underparts color (chin, throat, belly and vent combined). We standardized scores by adding values for each specimen and dividing by the sum of the highest value for each category (Toews et al. 2011; Céspedes-Arias et al. 2021; Aguillon and Rohwer 2022). Both facial and body hybrid indexes were used for cline fitting, with 0 representing typical *A. fuscoolivaceus* and 1 representing typical *A. flaviceps.* We excluded juveniles (identified by skull pneumatization, presence of Bursa of Fabricius, or juvenile plumage in photographs) but analyzed males and females together due to lack of sexual dichromatism.

### Cline fit analysis

We georeferenced all specimens and photographs and identified the northernmost record of typical *A. flaviceps* and the southernmost record of typical *A. fuscoolivaceus* (Fig. 1C). After tracing a straight line between these extreme endpoints, which spanned 573 km, we used this line as a geographic reference to assess phenotypic variation between species using cline fit analyses (Endler 1977; Barton and Hewitt 1985). We established the southern endpoint (*A. fuscoolivaceus*) as 0 km. We joined every locality point (from photographs and specimens) to the reference line We then projected every locality point perpendicular to the reference line and measured the distance from the southern endpoint to each projection point to determine the position of each specimen/photograph along the study transect.

For the facial hybrid index cline, we analyzed data from 111 photographs and 40 specimens, which we grouped into 16 sampling locations, and for the body color cline, we included 40 specimens grouped into 10 sampling locations. These locations were defined independently for each cline. We used the R package HZAR v.0.2.5 (Derryberry et al. 2014) to fit geographic clines along one-dimensional transects using the Metropolis-Hastings Markov chain Monte Carlo approximation. We ran HZAR using four equilibrium cline models (Szymura and Barton 1986, 1991; Gay et al. 2008). The best-fitting models for both traits provided estimates of cline width (w) and center (c), with fixed mean and variance for trait values at the transect extremes. We ran each model using a tuning value of 1.5 across three chains of 1 million steps, discarding the first 10% as burn-in. We assessed convergence through trace plots and randomized starting parameters for each run. We selected the best model using the Akaike Information Criterion with correction, AICc (DeRaad et al. 2019). Finally, we extracted the estimated cline center and width values with 95% Bayesian credibility intervals, and the maximum likelihood cline generated by the best model.

### Song discrimination analysis

Duets, songs, and calls of both species are extremely similar and essentially indistinguishable by ear. GMQ conducted fieldwork, with assistance from K. Certuche Cubillos, from 17–19 September 2022 in Juntas, above Ibagué, Tolima. This site, situated at 1825 meters above sea level on the eastern slope of the Central Cordillera, supports a relatively common population of our target species, *A. flaviceps*. Although we intended to conduct mirror experiment within the range of *A. fuscoolivaceus*, logistical and financial constraints prevented this. We followed the experimental design recommendations outlined in McGregor (2000). After identifying territorial pairs or groups suitable for playback trials, we exposed each subject to three pre-recorded stimuli (Fig. 2): a local song (*A. flaviceps,* based on XC531569), a foreign song (*A. fuscoolivaceus*, based on XC298289), and a control song (*Zonotrichia capensis,* based on XC693968.We broadcast each stimulus for 2 minutes using a JBL Clip 4 speaker, which we concealed in bushes one meter above the ground near the focal birds. We randomized the presentation order of foreign and control songs, while always presenting the local song last.

**Figure 2.**
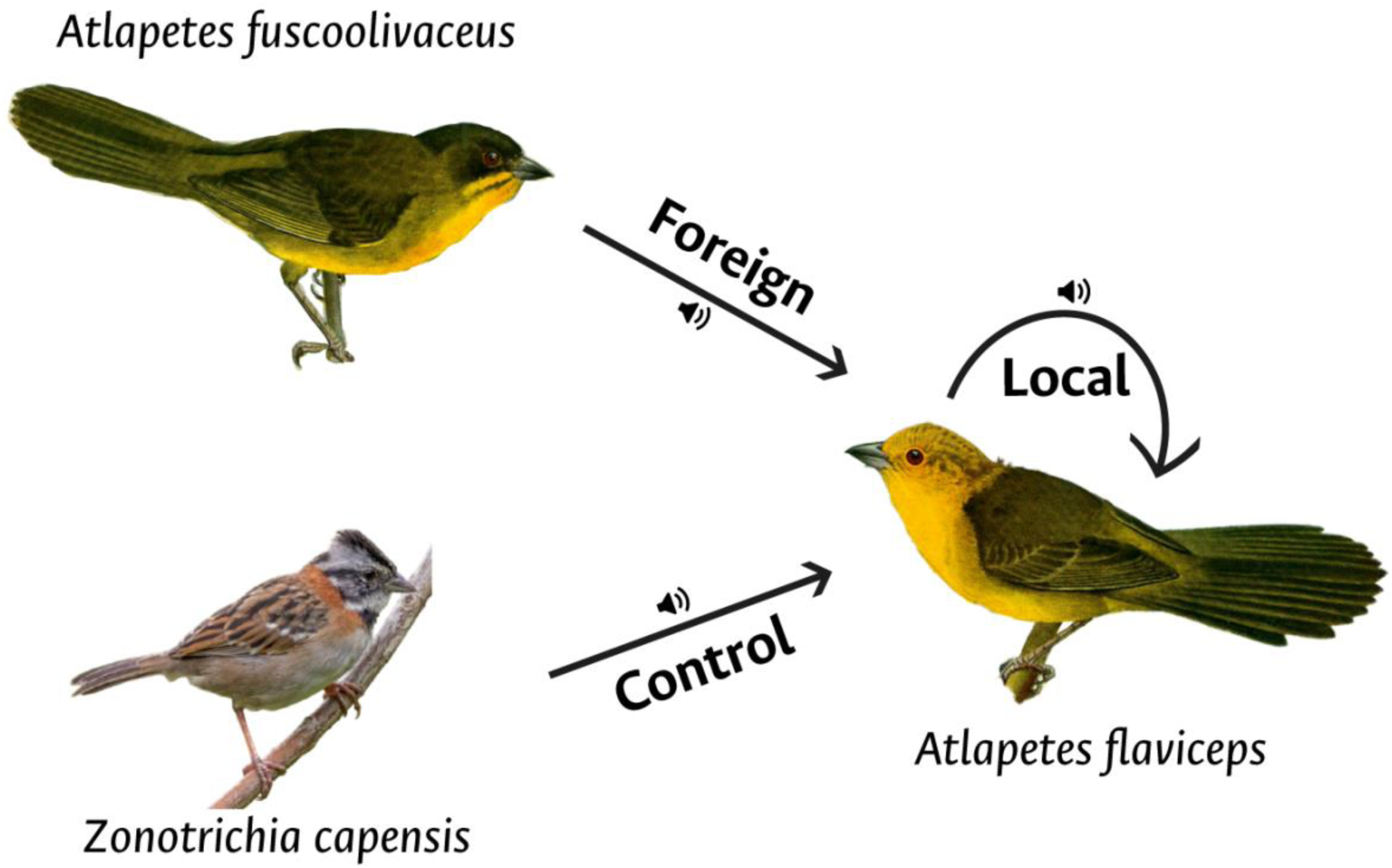
Scheme of the playback experiment design implemented in Juntas, Ibagué, Tolima, within the range of *Atlapetes flaviceps* (the local population). We reproduced either the foreign (*A. fuscoolivaceus*) or the control song (*Zonotrichia capensis*) first, followed by the playback of the local song (conspecific with *A. flaviceps*). Photo of *Z. capensis* by Charles Sharp.

We measured two behavioral variables to assess the subject’s responses to stimuli: latency and vocal display duration. Latency was defined as the time taken by individuals to begin responding to the stimulus, while duration was the time from the onset of the response until the focal bird ceased vocalizing and returned to their initial state. To analyze differences in response across treatments, we performed regression analyses in R (R Core Team 2021) using linear models, with song treatment (local, foreign, and control) as the fixed effect, and latency and duration as response variables. This allowed us to determine whether there were significant differences in responses to each song type. We then conducted a post hoc test using the *emmeans* 1.10.5 (Lenth 2016) to determine which treatments differed significantly from each other.

## RESULTS

In the contact zone between the ranges of pure *A. flaviceps* and *A. fuscoolivaceus*, we observed a remarkable diversity of facial and body coloration, resulting from unique combinations of color and pattern across individual plumage patches. As illustrated in Fig. 3, head coloration varies significantly among specimens, highlighting the extent of color variation beyond the extremes of each species’ range. We did not observe pure plumage forms in specimens or photographs of the two species together in the same locality. Instead, intermediate phenotypes were prevalent in southern Tolima, northern Huila, and eastern Cauca, exhibiting color combinations that diverge from the typical phenotypes of both species. For instance, intermediate individuals lack the bright yellow facial plumage of *A. flaviceps* and do not display the pronounced malar lines and dark facial features characteristic of *A. fuscoolivaceus*. Instead, they often exhibit greenish-olive tones throughout. Notably, an adult male *A. fuscoolivaceus* from the type locality in San Agustín, Huila (outside the contact zone) displayed scattered yellow feathers on the face, a rare feature not observed in other individuals from this population.

**Figure 3.**
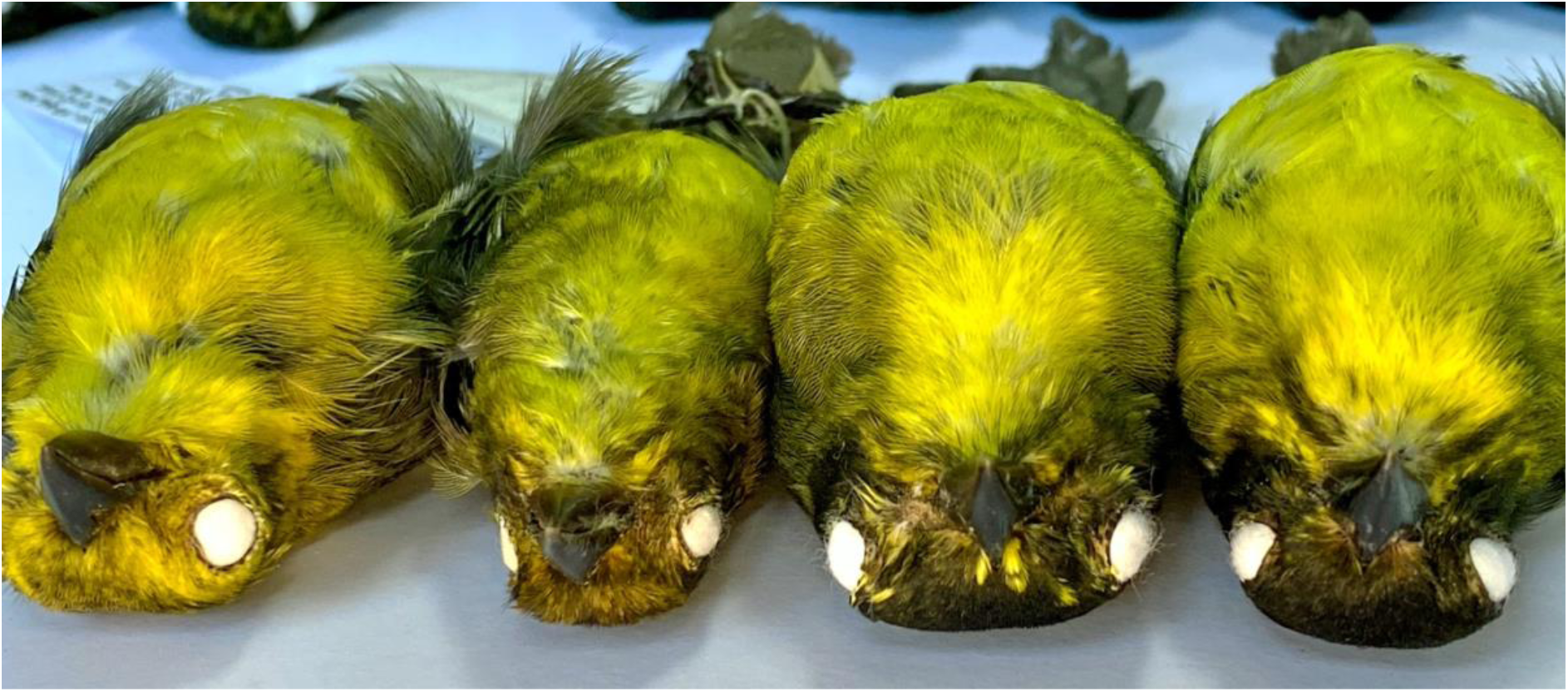
Examples of color and pattern variation in forehead and lores among selected specimens, from pure *A. fuscoolivaceus* (extreme left) to pure *A. flaviceps* (right). Photos by Colección Nacional de Aves, Instituto de Ciencias Naturales.

The two clines, of facial pattern and body coloration, exhibited sigmoidal shapes. The body coloration cline had a highly consistent center, at 132 km (near Inzá, Cauca), with an estimated width of 73 km (Fig. 4A). In comparison, the cline for facial pattern was centered at 131 km, near Inzá, Cauca, with a narrower width of 12 km (Fig. 4B).

**Figure 4.**
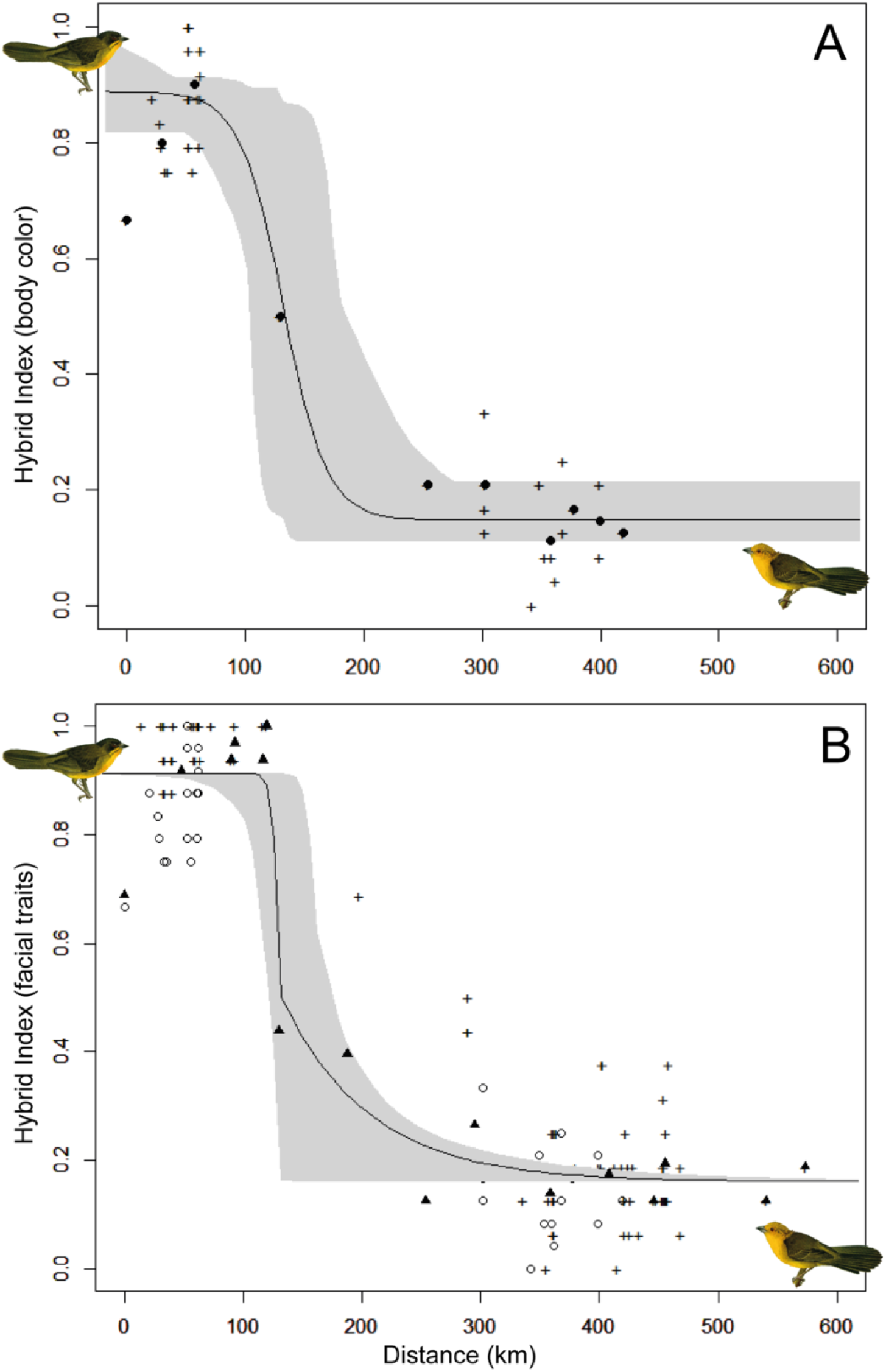
Geographic clines and associated 95% credibility intervals for two hybrid indexes, where higher values indicate typical *A. fuscoolivaceus* and lower values indicate typical *A. flaviceps.* **A)** facial color patterns, obtained from specimens and photographs (N=151) and based on four facial patches (Table 1). Black triangles denote mean values per locality, empty dots represent scores from specimens, and crosses from photographs. **B)** body plumage pattern and based on seven plumage patches scored from specimens (N=40). Crosses indicate mean hybrid index per locality, while black dots represent individual scores. Note the coincident cline center and much narrower width of the facial cline, and the need for denser sampling at the contact zone.

In total, we conducted playback experiments on seven *A. flaviceps* groups (six pairs and one group of five individuals). We found that *A. flaviceps* elicited an aggressive behavioral response equally to both its own song and the foreign song (i.e., that of *A. fuscoolivaceus*), but showed no response to the control. This response was initiated quickly, with an average latency of 0.27 minutes (Fig. 5A). Although there was no difference in latency between the local and foreign songs, response duration was significantly longer following the local song than the foreign stimulus (Fig. 5B).

**Figure 5.**
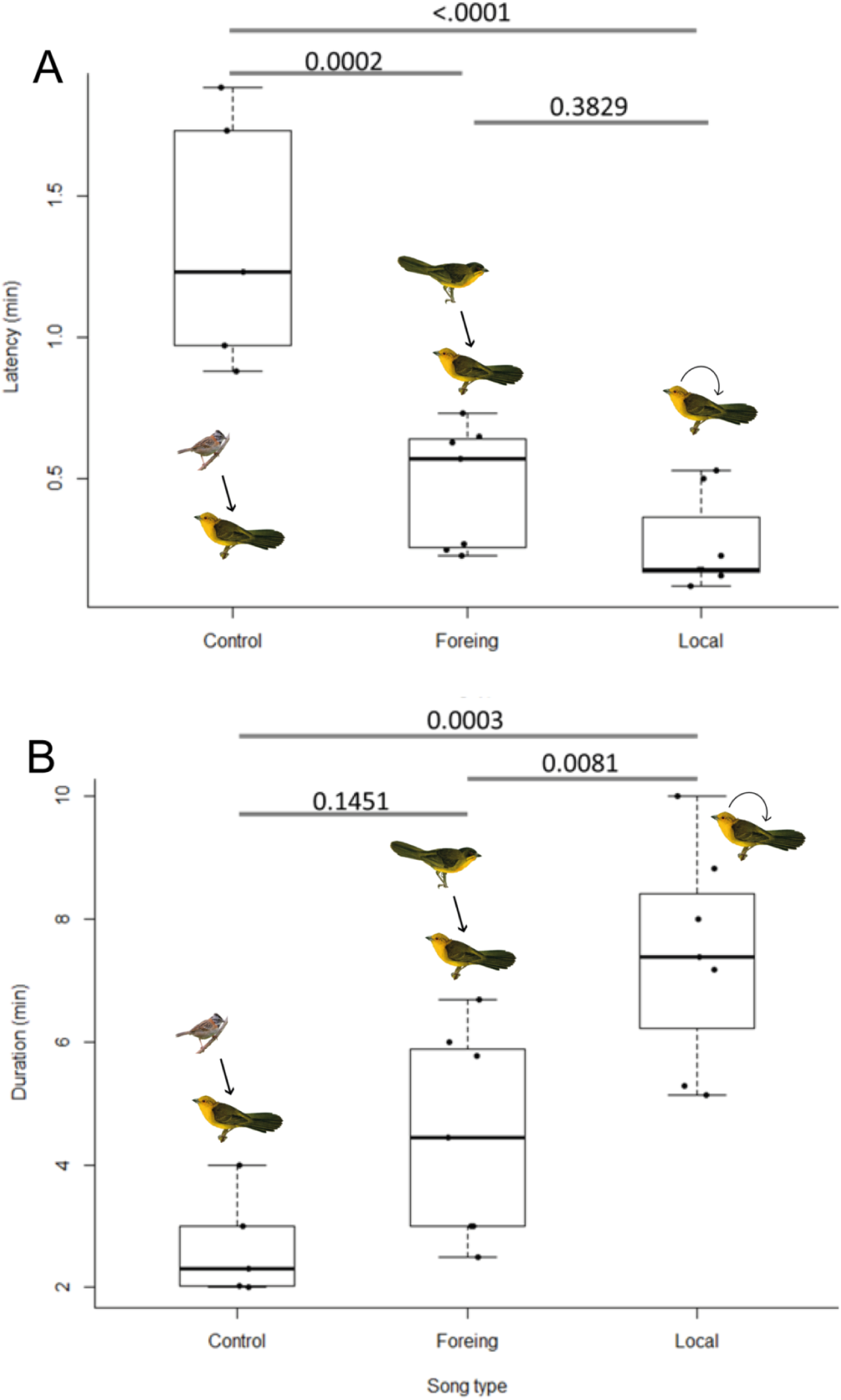
Vocal responses to playback stimuli. The top lines and numbers indicate *p*-values from post hoc tests. **A)** Latency of response for each song type. **B)** Duration of response for each song type. Local pairs or groups of *A. flaviceps* elicit an equally rapid response to its own (local) and to *A. fuscoolivaceus* (foreign) pre-recorded songs, but responses to the latter tended to last shorter than responses to local songs.

## DISCUSSION

In this study, we provide evidence for clinal phenotypic variation between *Atlapetes fuscoolivaceus* of the Upper Magdalena valley and *A. flaviceps* of the eastern slope of the middle Cordillera Central of Colombia. Accordingly, contrary to expectations of either a sharp replacement or sympatry without intermediates, we document a wide range of intermediate phenotypes at their presumed range boundary. This phenotypic variation, assessed along a geographic transect between the two forms, constructs a sigmoidal pattern that is predicted by hybridization (Endler 1977; Moore 1977). Thus, this clinal pattern reveals a novel, putative hybrid zone between these two brushfinches, whose limited behavioral discrimination further supports the potential for hybridization.

The putative hybrid zone, located in southern Tolima, northern Huila, and eastern Cauca, is formed by individuals with mixed plumage characteristics that diverge from the typical coloration of either species. Pure forms of *A. fuscoolivaceus* and *A. flaviceps* were only observed at the southern and northern extremes of the transect, respectively. Within the hybrid zone, however, we documented a spectrum of intermediate phenotypes, especially noticeable in the sides of head, lores, forehead, and crown, where color ranged from olive-yellow to dark olive-brown. Some individuals exhibited the yellow throat, underparts, and predominantly yellowish face of *A. flaviceps* but also showed a faint malar line, characteristic of *A. fuscoolivaceus*. Others had the darker head pattern and distinct malar line of *A. fuscoolivaceus*, though in light olive-green rather than the typical blackish hue, and with yellow lores and forehead. Malar lines also varied, ranging from wide and dark to thin and greenish across individuals. This variation has confused ornithologists and birdwatchers familiar with one of the species only. For instance, individuals photographed at La Argentina (formerly, La Plata Vieja, Huila) were identified as *A. flaviceps* (Dunning 1982) given its paler head and yellow lores and forehead that contrasted with the better-known darker *A. fuscoolivaceus* found 15 km apart at Reserva Natural Merenberg (Ridgely and Gaulin 1980). Interestingly, even *A. flaviceps* individuals outside the hybrid zone showed subtle color variation, with some showing hints of dark olive-green feathers on the face and crown. In addition, one adult male collected near the southern edge of the *A. fuscoolivaceus* range exhibited scattered bright yellow feathers within otherwise typical *A. fuscoolivaceus* plumage.

The geographic transition of phenotypic traits aligns with a classic hybrid zone pattern described by sigmoidal clines (Endler 1977; Moore and Price 1993). The centers of both body and facial color clines are located near Inzá, Cauca, but the cline widths differ significantly: facial coloration has a narrower while body coloration spans a broader width (12 vs. 73 km). This difference suggests stronger selection pressure on facial markings and patterns, which are the most distinct between the species, compared to overall body coloration, where both species are more similar. Stronger selection on facial pattern could enhance social or reproductive signaling when combined with gestures and vocalizations. However, differences in sample size may also influence this disparity, as facial pattern data were drawn from 151 individuals (40 specimens and 111 photographs), whereas body coloration data were limited to 40 specimens across a widwer geographic range. Increased sampling across the contact zone would help refine these estimates, providing more accurate cline width measures for body coloration. In any event, these findings suggest that hybridization may be driving the observed phenotypic diversity, with differential selection pressures likely acting on specific plumage traits.

The playback experiments further suggest the notion of weak reproductive isolation between *A. fuscoolivaceus* and *A. flaviceps*. In territories of *A. flaviceps*, birds responded aggressively to both conspecific (local) and heterospecific stimuli, showing no difference in response latency but a longer response duration to the local song. This subtle distinction may indicate a partial conspecific preference but suggests that vocal behavior alone may not fully prevent interbreeding in the contact zone. In other words, mechanisms of species recognition might not have evolved sufficiently to prevent behavioral interactions between the two species. Without strong vocal divergence, interbreeding may persist (Price 2008; Uy et al. 2018), contributing to gene flow and the maintenance of this hybrid zone.

The observed clinal variation in plumage and the lack of strong song discrimination align with hybrid zone dynamics typical of closely related species with overlapping ecological niches (Grant and Grant 1992; Moore and Price 1993; Price 2008; Uy et al. 2018). Interbreeding in birds can often occur despite some level of vocal divergence (Wilkins et al. 2013), especially when ecological differences are minimal. In our case, vocal divergence appears to be minor, although further quantitative analysis would be necessary to test that observation. Both species share similar habitats and elevational distributions along the eastern slope of the Central Cordillera and the Upper Magdalena valley, with their ranges merging geographically. In the contact zone, intermediate plumage patterns are common, while individuals showing typical parental forms are notably absent. This geographic overlap and the phenotypic sigmoidal gradient suggest a hybrid zone where ongoing gene flow in equilibrium with selection occurs within a narrow contact area.

Our findings highlight the role of hybridization in shaping biodiversity within the Colombian Andes (Morales-Rozo et al. 2017; Céspedes-Arias et al. 2021), where a diverse geological history, complex topography, climate fluctuations and vegetation shifts have driven pulses of isolation and connection. While studies of hybrid zones in the Neotropics remain limited (Ottenburghs 2023), our results add to the growing body of evidence suggesting that hybridization is a key process shaping this biodiversity hotspot (Morales-Rozo et al. 2017; Pulido-Santacruz et al. 2018; Céspedes-Arias et al. 2021; Del-Rio et al. 2022). In the Andes, geographic or ecological barriers may only partially restrict gene flow between recently diverged lineages (Prieto-Torres et al. 2018; Palacios et al. 2019). This initial evaluation of a new hybrid zone in the Andes lays the groundwork for long-term studies on the *Atlapetes* species pair as a valuable model for investigating speciation and hybridization in tropical montane birds.

### Future directions

To deepen our understanding of this hybrid zone, we recommend expanding the sampling transect of specimens across the entire range of each species. We are developing a new study, led by K. Certuche Cubillos, that aims to fill current sampling gaps, especially along the hybrid zone between La Plata, Huila, northward to Inzá, Cauca, and the slopes of Nevado del Huila, as well as further north to Planadas and Chaparral, Tolima. We plan to sample the isolated pure populations of *A. flaviceps* in the northern sector the in Antioquia, Caldas or Risaralda (Calderón-Franco et al. 2012; López-Ordóñez et al. 2013; Chaparro-Herrera et al. 2020), and the isolated populations of *A. fuscoolivaceus* in the Eastern Cordillera near Alpujarra, Tolima and Colombia, Huila, and the eastern Amazonian slope in Cauca (Salaman et al. 2002; Gallo-Cajiao et al. 2014; Gómez-Bernal et al. 2016; Ruiz-Ovalle and de Oliveira 2020). These new locations would expand the cline’s length and further refine its estimated parameters. While we expect the center near Inzá will remain consistent, additional data may lead to narrower and more reliable confidence intervals for phenotypic cline widths. The expanded sampling will allow us to assess morphological variation through morphometric clines, quantify coloration using spectrographic data, in addition to updating the phenotypic scores of facial and body coloration presented herein.

In parallel, we are sequencing an annotated, chromosome-level reference genome of a pure *A. fuscoolivaceus* individual from San Agustín, Huila. This genomic resource will enable a wide range of genetic analyses of the hybrid zone using our vouchered tissue samples, including assessments of population structure and historical demography, genetic introgression, identification of loci under selection, gene regions associated with the variable facial traits, and the extent of selection against hybrids. Integrating these genetic analyses will be crucial to evaluate the origin of the hybrid zone and the timing of speciation in *Atlapetes*.

Additional ecological and behavioral studies would shed light on the mechanisms of divergence and hybridization. Our exploratory one-sided playback experiments suggest that *A. flaviceps* individuals do not strongly discriminate against *A. fuscoolivaceus* songs, though their response was less persistent than to conspecific songs. Additional replicate assays, fine-tuned measurements of response intensity, and reciprocal studies in the range of pure *A. fuscoolivaceus* would help clarify the role of vocalizations and behavior in maintaining or eroding species boundaries. Together, these steps would provide deeper insights into the dynamics of this hybrid zone and the evolutionary processes driving speciation of *Atlapetes* in the Andes.

## ACKNOWLEDGMENTS

This manuscript is the result of GMQ’s undergraduate thesis at the National University of Colombia, inspired by our recent observations of intermediate individuals in Huila and Tolima. We gratefully acknowledge our friends and colleagues from the Colombia Resurvey Project, especially Natalia Pérez, Juliana Soto, Nelsy Niño, Jessica Díaz, Andrés Sierra, Glenn Seeholzer, Natalia Ocampo, Daniel Cadena, and Camila Gómez, as well as participants in the “Alas, Cantos y Colores” expeditions, funded by the Minciencias Colombia BIO program and organized by the Alexander von Humboldt Institute and the Institute of Natural Sciences, National University of Colombia (ICN). We are deeply grateful to Camilo Váquiro, Sebastián Betancourt, Daniela Pulido, Rosalino Ortiz, and Sergio Barreiro, whose support was essential to our fieldwork in Huila. Special thanks to bird curators Sergio Lozada at the Zoology Collection, University of Tolima (CZUT-OR), Luis Germán Gómez at the Museum of Natural History, University of Cauca (MHN-UC-AV), and Gustavo Bravo at the Alexander von Humboldt Institute (IAvH-A) for facilitating our visits and help with specimen loans. Ronald Fernández provided valuable guidance for the playback experiments, and Diego Cueva offered insightful discussion on hybridization. We appreciate the recommendations and mentorship of Jorge Pérez-Emán. We are also thankful to Katherine Certuche Cubillos for her support and companionship in the field during playback experiments in *A. flaviceps* territories. Finally, the feedback and encouragement of the ORNIS research group, particularly Esteban Lara, greatly enriched this study. We appreciate the contribution of georeferenced photographs of birdwatchers to public databases such as Macaulay Library via ebird.org and iNaturalist.

## Supplementary information

**Table S1.** List of specimens examined, and scored for both facial and body plumage patterns. The taxon identification as provided in the specimen labels, is binary. Those with asterisks exhibited an intermediate phenotype, and are from the putative hybrid zone area.

| Specimen number | Taxon ID* | Department | Locality | Latitude | Longitud |
| --- | --- | --- | --- | --- | --- |
| IAvH-A-7870 | *A. fuscoolivaceus* | Huila | Cueva de los Guácharos, Cabaña "La Iascoa" | 1.36443 | -76.22473 |
| MHNUC-AV-4591 | *A. fuscoolivaceus* | Cauca | Santa Rosa, San Juan de Villalobos | 1.55519 | -76.30527 |
| IAvH-A-11782 | *A. fuscoolivaceus* | Huila | Acevedo, PNN Cueva de los Guácharos | 1.61638 | -76.10436 |
| IAvH-A-11801 | *A. fuscoolivaceus* | Huila | Acevedo, PNN Cueva de los Guácharos | 1.62261 | -76.10452 |
| IAvH-A-13921 | *A. fuscoolivaceus* | Huila | Palestina, vda La Guajira, Reserva Natural La Riviera | 1.65638 | -76.18694 |
| IAvH-A-14102 | *A. fuscoolivaceus* | Huila | Palestina | 1.67444 | -76.07388 |
| ICN-42265 | *A. fuscoolivaceus* | Huila | San Agustín | 1.83061 | -76.35334 |
| ICN-42331 | *A. fuscoolivaceus* | Huila | San Agustín | 1.83122 | -76.35294 |
| ICN-42215 | *A. fuscoolivaceus* | Huila | San Agustín | 1.83171 | -76.35059 |
| ICN-42120 | *A. fuscoolivaceus* | Huila | San Agustín | 1.83171 | -76.35059 |
| ICN-42297 | *A. fuscoolivaceus* | Huila | San Agustín | 1.83177 | -76.35185 |
| ICN-42301 | *A. fuscoolivaceus* | Huila | San Agustín | 1.83184 | -76.35193 |
| ICN-41944 | *A. fuscoolivaceus* | Huila | San Agustín | 1.83216 | -76.35215 |
| ICN-42102 | *A. fuscoolivaceus* | Huila | San Agustín | 1.83257 | -76.35426 |
| ICN-3169 | *A. fuscoolivaceus* | Huila | San Agustín, La Candela | 1.863876 | -76.361492 |
| ICN-42375 | *A. fuscoolivaceus* | Huila | San Agustín | 1.90279 | -76.3787 |
| ICN-42376 | *A. fuscoolivaceus* | Huila | San Agustín | 1.90291 | -76.37864 |
| ICN-42519 | *A. fuscoolivaceus* | Huila | San Agustín | 1.91373 | -76.29552 |
| ICN-42510 | *A. fuscoolivaceus* | Huila | San Agustín | 1.91442 | -76.2931 |
| ICN-42488 | *A. fuscoolivaceus* | Huila | San Agustín | 1.91442 | -76.2931 |
| ICN-41946 | *A. fuscoolivaceus* | Huila | San Agustín | 1.91442 | -76.2931 |
| ICN-27333* | *A. fuscoolivaceus* | Huila | Inzá, vda. Tierras Blancas, vía Inzá-Gabriel López | 2.515896 | -76.077571 |
| ICN-27336* | *A. fuscoolivaceus* | Huila | Inzá, vda. Tierras Blancas, vía Inzá-Gabriel López | 2.515896 | -76.077571 |
| ICN-27370* | *A. fuscoolivaceus* | Huila | Inzá, Rio Guanacas, vda. Tierras Blancas, vía Popayán-Inzá | 2.525158 | -76.077931 |
| CZUT-0798 | *A. flaviceps* | Tolima | Río Blanco, Las Delicias, Bocatoma | 3.60716 | -75.65302 |
| CZUT-0996* | *A. flaviceps* | Tolima | Roncesvalles, vda. San Marcos | 4.03105 | -75.57458 |
| CZUT-1081* | *A. flaviceps* | Tolima | Roncesvalles, vda. El Diamante | 4.03333 | -75.56169 |
| CZUT-1079* | *A. flaviceps* | Tolima | Roncesvalles, vda. El Diamante | 4.03333 | -75.56169 |
| CZUT-1078* | *A. flaviceps* | Tolima | Roncesvalles, vda. El Diamante | 4.03333 | -75.56169 |
| CZUT-0053 | *A. flaviceps* | Tolima | Cajamarca, Carrizales | 4.38861 | -75.51388 |
| ICN-38782 | *A. flaviceps* | Tolima | Cajamarca, vda. La Luisa, Quebrada La Colosa, Sector La Vara | 4.446995 | -75.42468 |
| CZUT-1630 | *A. flaviceps* | Tolima | Ibagué, Coello San Juan Machin, Las brizas | 4.48102 | -75.39416 |
| IAvH-A-17676 | *A. flaviceps* | Tolima | Ibagué, Finca La Estrella | 4.53012 | -75.40528 |
| IAvH-A-17731 | *A. flaviceps* | Tolima | Ibagué, Finca La Estrella | 4.54988 | -75.46809 |
| IAvH-A-17730 | *A. flaviceps* | Tolima | Ibagué, Finca La Estrella | 4.54988 | -75.46809 |
| CZUT-1366 | *A. flaviceps* | Tolima | Ibagué, Juntas, Finca Ukukú | 4.55836 | -75.32777 |
| CZUT-0485 | *A. flaviceps* | Tolima | Santa Isabel, Guimaral | 4.69391 | -75.10558 |
| CZUT-0602 | *A. flaviceps* | Tolima | Líbano, El agrado | 4.88675 | -75.11072 |
| CZUT-0572 | *A. flaviceps* | Tolima | Líbano, El Agrado | 4.88675 | -75.11072 |
| CZUT-0601 | *A. flaviceps* | Tolima | Casabianca, Palma Peñitas | 5.06969 | -75.10161 |

**Table S2.** List of photographs examined and scored to calculate the facial hybrid index. The taxon identification as provided in the original source, is binary. Those with asterisks exhibited an intermediate phenotype, and are from the proposed hybrid zone area.

| Photo ID | Source | Taxon | Department | Locality | Latitude | Longitud |
| --- | --- | --- | --- | --- | --- | --- |
| P95 | ML433029981 | *A. fuscoolivaceus* | Huila | Vía Pitalito - Mocoa | 1.4933104 | -76.405429 |
| P111 | ML60833341 | *A. fuscoolivaceus* | Huila | Vía Mocoa-Pitalito, escuela abandonada | 1.6431535 | -76.258432 |
| P104 | ML321506501 | *A. fuscoolivaceus* | Huila | Sendero Los Chorros | 1.6583791 | -76.02345 |
| P86 | ML412637601 | *A. fuscoolivaceus* | Huila | Palestina, Reserva La Drymophila | 1.6608855 | -76.109527 |
| P87 | ML203311391 | *A. fuscoolivaceus* | Huila | Palestina, Reserva La Drymophila | 1.6608855 | -76.109527 |
| P88 | ML393378491 | *A. fuscoolivaceus* | Huila | Palestina, Reserva La Drymophila | 1.6608855 | -76.109527 |
| P98 | ML312156231 | *A. fuscoolivaceus* | Huila | Palestina, Reserva La Drymophila | 1.6608855 | -76.109527 |
| P108 | ML384398281 | *A. fuscoolivaceus* | Huila | Pitalito, Base Militar El Cable | 1.6662952 | -76.229779 |
| P107 | ML183881331 | *A. fuscoolivaceus* | Huila | Palestina, Reserva La Drymophila | 1.667379 | -76.106996 |
| P78 | ML142754031 | *A. fuscoolivaceus* | Huila | Palestina, Reserva El Encanto | 1.7201319 | -76.119332 |
| P79 | ML57741331 | *A. fuscoolivaceus* | Huila | Palestina, Reserva El Encanto | 1.7201319 | -76.119332 |
| P80 | ML57741311 | *A. fuscoolivaceus* | Huila | Palestina, Reserva El Encanto | 1.7201319 | -76.119332 |
| P81 | ML53629411 | *A. fuscoolivaceus* | Huila | Palestina, Reserva El Encanto | 1.7201319 | -76.119332 |
| P102 | ML178040901 | *A. fuscoolivaceus* | Huila | Reserva Natural Los Ariscos | 1.7328076 | -76.22352 |
| P76 | ML376516491 | *A. fuscoolivaceus* | Huila | Alto Santa Bárbara | 1.8670787 | -75.947747 |
| P91 | ML216081561 | *A. fuscoolivaceus* | Huila | Parque Arqueológico de San < | 1.8872471 | -76.295157 |
| P92 | ML216081671 | *A. fuscoolivaceus* | Huila | Parque Arqueológico de San Agustín | 1.8872471 | -76.295157 |
| P110 | ML139448701 | *A. fuscoolivaceus* | Huila | Cañón de Quebrada Negra | 1.899774 | -76.424928 |
| P106 | ML61956631 | *A. fuscoolivaceus* | Huila | Reserva Natural Monte Bonito | 1.9128321 | -76.096732 |
| P89 | ML24186811 | *A. fuscoolivaceus* | Huila | Parque Ecológico Alto de los Ídolos | 1.9160704 | -76.240643 |
| P90 | ML24090581 | *A. fuscoolivaceus* | Huila | Parque Ecológico Alto de los Ídolos | 1.9160704 | -76.240643 |
| P82 | ML256034421 | *A. fuscoolivaceus* | Huila | Finca La Guajira | 1.9180514 | -76.212785 |
| P83 | ML256034461 | *A. fuscoolivaceus* | Huila | Finca La Guajira | 1.9180514 | -76.212785 |
| P109 | ML239385981 | *A. fuscoolivaceus* | Huila | Reserva Natural Finca El Manantial | 1.9336819 | -76.229538 |
| P84 | ML216081431 | *A. fuscoolivaceus* | Huila | Bordones | 2.0102387 | -76.164942 |
| P105 | ML226788191 | *A. fuscoolivaceus* | Huila | Garzón, Vereda Las Mercedes | 2.1588954 | -75.561848 |
| P93 | ML56560381 | *A. fuscoolivaceus* | Huila | Parque Regional Serranía de Minas | 2.1871212 | -75.93956 |
| P94 | ML88718851 | *A. fuscoolivaceus* | Huila | Parque Regional Serranía de Minas | 2.1871212 | -75.93956 |
| P85 | ML180562511* | *A. fuscoolivaceus* | Huila | San Andrés, La Plata | 2.39158 | -75.838358 |
| P75 | ML170382081* | *A. fuscoolivaceus* | Huila | La Plata | 2.3965919 | -75.828724 |
| P97 | ML416130551* | *A. fuscoolivaceus* | Huila | Reserva Ecoturística El Carriquí y el Barranquero | 2.426611 | -75.460639 |
| P49 | ML116518601* | *A. flaviceps* | Huila | Santa María, Vereda San Francisco | 2.975452 | -75.656457 |
| P29 | ML111035181* | *A. flaviceps* | Huila | PNN Nevado del Huila, Sector el Roble | 2.9900131 | -75.67061 |
| P96 | ML97066601* | *A. fuscoolivaceus* | Tolima | Planadas, Bosque Cami | 3.1159863 | -75.633747 |
| P3 | ML459305281* | *A. flaviceps* | Tolima | Ortega, Camino Real | 3.9173288 | -75.454568 |
| P4 | ML310198231* | *A. flaviceps* | Tolima | Ortega, Camino Real | 3.9173288 | -75.454568 |
| P99 | ML251428091* | *A. fuscoolivaceus* | Tolima | Reserva Natural Finca El Manantial | 3.9271623 | -75.444419 |
| P100 | ML251428081* | *A. fuscoolivaceus* | Tolima | Reserva Natural Finca El Manantial | 3.9271623 | -75.444419 |
| P101 | ML251428101* | *A. fuscoolivaceus* | Tolima | Reserva Natural Finca El Manantial | 3.9271623 | -75.444419 |
| P17 | ML437700581 | *A. flaviceps* | Tolima | Cajamarca, Finca La Tribuna | 4.333 | -75.515 |
| P65 | ML346155101 | *A. flaviceps* | Tolima | Via Toche | 4.4506692 | -75.430158 |
| P66 | ML321925471 | *A. flaviceps* | Tolima | RN Valle Escondido | 4.5001949 | -75.294207 |
| P67 | ML314774081 | *A. flaviceps* | Tolima | Parque Ecoturístico La Plata | 4.5204308 | -75.287822 |
| P42 | ML466950271 | *A. flaviceps* | Tolima | Río Toche | 4.544855 | -75.408766 |
| P8 | ML117624331 | *A. flaviceps* | Tolima | Ibagué, Cañón del Combeima, Juntas | 4.5546418 | -75.322394 |
| P9 | ML368286041 | *A. flaviceps* | Tolima | Ibagué, Cañón del Combeima, Juntas | 4.5546418 | -75.322394 |
| P10 | ML71012261 | *A. flaviceps* | Tolima | Ibagué, Cañón del Combeima, Juntas | 4.5546418 | -75.322394 |
| P44 | Private | *A. flaviceps* | Tolima | Ukuku Rural Lodge | 4.5582438 | -75.327265 |
| P45 | ML23771021 | *A. flaviceps* | Tolima | Ukuku Rural Lodge | 4.5582438 | -75.327265 |
| P46 | ML361928951 | *A. flaviceps* | Tolima | Ukuku Rural Lodge | 4.5582438 | -75.327265 |
| P47 | ML411745941 | *A. flaviceps* | Tolima | Ukuku Rural Lodge | 4.5582438 | -75.327265 |
| P54 | ML470529311 | *A. flaviceps* | Tolima | Ukuku Rural Lodge | 4.5582438 | -75.327265 |
| P55 | ML469949621 | *A. flaviceps* | Tolima | Ukuku Rural Lodge | 4.5582438 | -75.327265 |
| P59 | ML435695921 | *A. flaviceps* | Tolima | Ukuku Rural Lodge | 4.5582438 | -75.327265 |
| P62 | ML396456381 | *A. flaviceps* | Tolima | Ukuku Rural Lodge | 4.5582438 | -75.327265 |
| P73 | ML268427531 | *A. flaviceps* | Tolima | Ukuku Rural Lodge | 4.5582438 | -75.327265 |
| P57 | ML457349871 | *A. flaviceps* | Tolima | Ukuku Rural Lodge | 4.5586359 | -75.327917 |
| P5 | Private | *A. flaviceps* | Tolima | Ibagué, Cañón del Combeima | 4.559751 | -75.320791 |
| P6 | ML195067861 | *A. flaviceps* | Tolima | Ibagué, Cañón del Combeima, Cabaña Dulima | 4.5786066 | -75.324929 |
| P7 | ML195069011 | *A. flaviceps* | Tolima | Ibagué, Cañón del Combeima, Cabaña Dulima | 4.5786066 | -75.324929 |
| P74 | ML195069041 | *A. flaviceps* | Tolima | Cañón del Río Combeima, Cabaña Dulima | 4.5786066 | -75.324929 |
| P13 | ML358350141 | *A. flaviceps* | Tolima | Santa Isabel, Via El bosque | 4.7142617 | -75.105087 |
| P68 | ML308488221 | *A. flaviceps* | Tolima | Camino La Tigrera | 4.9014448 | -75.082256 |
| P56 | ML458775201 | *A. flaviceps* | Tolima | Hacienda Los Tejos | 4.9166024 | -75.076093 |
| P14 | ML344043381 | *A. flaviceps* | Tolima | El Líbano | 4.9235085 | -75.081474 |
| P15 | ML344043401 | *A. flaviceps* | Tolima | El Líbano | 4.9235085 | -75.081474 |
| P16 | ML344043331 | *A. flaviceps* | Tolima | El Líbano | 4.9235085 | -75.081474 |
| P40 | ML391279771 | *A. flaviceps* | Tolima | Villahermosa | 5.0236955 | -75.045914 |
| P53 | ML327689041 | *A. flaviceps* | Tolima | Villahermosa y alrededores | 5.0298527 | -75.122549 |
| P23 | ML401745131 | *A. flaviceps* | Tolima | Herveo, Miraderos | 5.0649574 | -75.203526 |
| P19 | ML118162331 | *A. flaviceps* | Tolima | Herveo | 5.0799273 | -75.177606 |
| P37 | ML457942181 | *A. flaviceps* | Tolima | Sector Numero 6 | 5.0860928 | -75.171438 |
| P38 | ML457942411 | *A. flaviceps* | Tolima | Sector Numero 6 | 5.0860928 | -75.171438 |
| P50 | ML450044951 | *A. flaviceps* | Tolima | Herveo, Vereda Torreseis | 5.0932483 | -75.171098 |
| P34 | ML430745791 | *A. flaviceps* | Risaralda | Apía | 5.1144907 | -75.9286 |
| P1 | ML129774511 | *A. flaviceps* | Risaralda | Apia, Aguabonita | 5.1220625 | -75.942392 |
| P41 | ML400403761 | *A. flaviceps* | Tolima | Padua | 5.1278911 | -75.148667 |
| P28 | ML207388071 | *A. flaviceps* | Risaralda | Pirineos Ecolodge | 5.1421813 | -75.948878 |
| P70 | ML288962611 | *A. flaviceps* | Risaralda | PN Santa Emilia | 5.2035273 | -75.901237 |
| P39 | ML28778191 | *A. flaviceps* | Risaralda | Sendero la Linda, Vereda La Aldea | 5.3039878 | -75.993106 |
| P61 | ML407730131 | *A. flaviceps* | Risaralda | San Clemente, La Ceiba | 5.3222118 | -75.7818 |
| P30 | ML331025591 | *A. flaviceps* | Tolima | Reserva El Popal | 5.3710396 | -75.186095 |
| P32 | ML414917441 | *A. flaviceps* | Caldas | Río Pensilvania | 5.37591 | -75.169235 |
| P33 | ML414917461 | *A. flaviceps* | Caldas | Río Pensilvania, Confa | 5.37591 | -75.169235 |
| P71 | ML281952101 | *A. flaviceps* | Caldas | El Bosque | 5.3760978 | -75.169646 |
| P11 | ML238943221 | *A. flaviceps* | Tolima | El Bosque | 5.3777527 | -75.169788 |
| P12 | ML238943261 | *A. flaviceps* | Tolima | El Bosque | 5.3777527 | -75.169788 |
| P27 | ML347243511 | *A. flaviceps* | Caldas | Pensilvania | 5.385785 | -75.165164 |
| P31 | ML402474671 | *A. flaviceps* | Caldas | Ríosucio, Reserva Lifer | 5.39467 | -75.705507 |
| P58 | ML445446161 | *A. flaviceps* | Tolima | Pensilvania, Cascadas, La Arenera | 5.3956081 | -75.156367 |
| P63 | ML393635141 | *A. flaviceps* | Tolima | Pensilvania, Cascadas, La Arenera | 5.3956081 | -75.156367 |
| P72 | ML278481631 | *A. flaviceps* | Caldas | Pensilvania, Repetidora | 5.4030735 | -75.149072 |
| P64 | ML380026111 | *A. flaviceps* | Risaralda | Carretera a Río Sucio | 5.4035916 | -75.714494 |
| P60 | ML431751561 | *A. flaviceps* | Caldas | Pensilvania, La Repetidora | 5.4077778 | -75.146944 |
| P2 | ML348475001 | *A. flaviceps* | Caldas | Aldea Sol de los Andes | 5.4973 | -75.68348 |
| P69 | ML294798541 | *A. flaviceps* | Caldas | Bermajal, Resguardo San Lorenzo | 5.5083401 | -75.717072 |
| P52 | ML156494101 | *A. flaviceps* | Antioquia | Vía Parque Caldas-Angelópolis | 6.133813 | -75.679021 |
| P48 | ML93520151 | *A. flaviceps* | Antioquia | Vereda La Chapa | 6.4226924 | -75.281137 |

## Notes

### Competing Interest Statement

The authors have declared no competing interest.

